# A high-quality *de novo* genome assembly from a single parasitoid wasp

**DOI:** 10.1101/2020.07.13.200725

**Authors:** Xinhai Ye, Yi Yang, Zhaoyang Tian, Le Xu, Kaili Yu, Shan Xiao, Chuanlin Yin, Shijiao Xiong, Qi Fang, Hu Chen, Fei Li, Gongyin Ye

## Abstract

Sequencing and assembling a genome with a single individual have several advantages, such as lower heterozygosity and easier sample preparation. However, the amount of genomic DNA of some small sized organisms might not meet the standard DNA input requirement for current sequencing pipelines. Although few studies sequenced a single small insect with about 100 ng DNA as input, it may still be challenging for many small organisms to obtain such amount of DNA from a single individual. Here, we use 20 ng DNA as input, and present a high-quality genome assembly for a single haploid male parasitoid wasp (*Habrobracon hebetor*) using Nanopore and Illumina. Because of the low input DNA, a whole genome amplification (WGA) method is used before sequencing. The assembled genome size is 131.6 Mb with a contig N50 of 1.63 Mb. A total of 99% Benchmarking Universal Single-Copy Orthologs are detected, suggesting the high level of completeness of the genome assembly. Genome comparison between *H. hebetor* and its relative *Bracon brevicornis* shows a high-level genome synteny, indicating the genome of *H. hebetor* is highly accurate and contiguous. Our study provides an example for *de novo* assembling a genome from ultra-low input DNA, and will be used for sequencing projects of small sized species and rare samples, haploid genomics as well as population genetics of small sized species.

## Introduction

A high-quality genome assembly is one of the most important resources for studying biological questions in organisms. However, genome sequencing and assembly can be complicated by the small body size of many organisms (i.e., very low genomic DNA from a single individual) and high heterozygosity [1, 2]. In particular, many arthropod (insect) genome projects face these problems, and obtain highly fragmented genomes with very low Contig N50 or/and Scaffold N50 value [1]. These fragmented genomes will have problems in genome annotation, as some genes are incomplete in genome assembly. In addition, gene synteny analysis, chromosome evolution, quantitative trait locus mapping also will fail in such fragmented genomes [1].

Mostly, the amount of genomic DNA is very low when obtained from single small-sized insect, which makes it hard to meet the standard DNA input requirement of long-reads sequencing (Pacbio or Nanopore) or even short-reads sequencing (Illumina) [1, 3–6]. Over the past two decades, many insects with small body size (e. g., parasitoid wasps, aphids, many *Drosophila*) were sequenced by using the DNA from pooled samples [2]. But pooling method raises heterozygosities in genomic regions, which will be assembled into more fragmented contigs. To reduce the heterozygosity level in pooled sample, inbreeding species were used for DNA extraction and sequencing in many cases [1, 7, 8]. Using the current hybrid genome sequencing and assembly approaches, many high-quality genome assemblies (some are chromosome-level genomes) were released [9]. However, most of insects are difficult to collect or cannot be well reared in the lab. Even if they can be reared in the lab, they might be difficult to inbreed [1, 3, 6]. Therefore, obtaining a high-quality genome assembly is still a problem for some small sized species and rare species in the wild.

Recently, to resolve these problems, some approaches were developed in sequencing from a single individual with low DNA input. Kingan et al. reported a genome assembly from a single mosquito (about 100 ng genomic DNA), *Anopheles coluzzii*, sequenced with three PacBio SMRT Cells [6]. Adams et al. developed a hybrid method (Illumina, Nanopore and Hi-C) to obtain a chromosome-level genome assembly from a single *Drosophila melanogaster* (totally ~200ng genomic DNA) [10]. These two studies provided good examples for sequencing a single small insect, but obtaining about 100 ng DNA from a single individual may still be challenging for many small insects such as parasitoid wasps.

How to sequence and assemble a high-quality genome from a single insect with ultra-low input DNA is still a problem, and no practical experience in this field up to now. Because of the ultra-low input DNA, it is difficult to construct library for sequencing at this time. A whole genome amplification (WGA) method has to be used to increase the total amount of DNA to meet the lowest requirement of sequencing library construction [11]. The WGA method is widely used to identify single-nucleotide polymorphisms, copy number variations in low DNA sample, e. g. single cell [12, 13]. However, this method is rarely used in *de novo* genome assembly. It is important to note that WGA has some disadvantages, such as potential amplification biases and contaminant problems [12, 14–16], which might influence the quality of genome assembly. There are only a few cases on *de novo* genome assembly based on WGA data, most of them are in bacteria [17, 18]. Recently, WGA was used to build a *de novo* genome assembly for a fungus [19]. But there is no report about WGA application in *de novo* genome assembly in insects or other more complex eukaryotic species.

Here, starting with 20 ng DNA, we present a high-quality *de novo* genome assembly of a single male parasitoid wasp (*Habrobracon hebetor*) using WGA and Nanopore, Illumina sequencing. This approach provides an example for genome sequencing and assembly using ultra-low DNA input, and is applicable for small size organisms and rare samples.

## Results and discussion

### Parasitoid wasp for sequencing

Parasitoid wasps are interesting and important organisms for studying fundamental biological questions such as evolution and sex determination, and some of them are important natural enemies for insect pest management [20, 21]. In addition, parasitoid wasps are often in very small size, which makes the genome projects of them complicated. Although many studies have assembled genomes from pooled inbred lab strains [8, 22–27], most of parasitoid wasps are difficult to be reared in the lab to establish such lab strains for sequencing. There are also some problems and uncertain factors in sequencing field collected samples, such as high heterozygosity and insufficient sample. These issues are major hindrances to the development of parasitoid wasp genomics. To test the feasibility of sequencing and assembling genome from a single parasitoid wasp, we sequenced a single male adult wasp of *H. hebetor* (Braconidae) in the study. *H. hebetor* wasp is an important biological control agent for managing multiple lepidopteran pests [28], and an ideal model for Hymenoptera sex determination researches [29, 30].

### Genome sequencing

In total, 102 ng high molecular weight DNA was extracted from a single male adult wasp of *H. hebetor*. Two studies already had provided good cases to obtain a high-quality genome assembly with approximately 100 ng of DNA [6, 10]. However, it still might be difficult to obtain ~100 ng DNA from a single individual in many small sized species. To challenge the lower input DNA for sequencing and make our method useful for more small size species, only 20 ng DNA was used for subsequent study.

DNA was subjected to whole genome amplification, yielding ~4.37 μg amplified DNA. 2.02 μg amplified DNA was used for Nanopore sequencing. In total, we obtained 51 Gb high-quality reads (~372X, genome size is about 137 Mb) from a single Oxford Nanopore Technology (ONT) PromethION flow cell (Table. S1). According to the handbook of this WGA method, the average amplified product length is in a range between 2 kb and 100 kb. The distribution of the Nanopore reads showed the similar pattern (Table. S1 and Figure. S1). The average length of total reads is 4,484 bp, and the N50 of total reads is 7,311 bp. We also generated 13.8 Gb Illumina clean data reads using the rest of amplified DNA (Table. S2).

### Genome assembly

We assembled a genome assembly using Flye [31, 32] with ~250X >5K Nanopore reads. This assembly was then corrected and polished by both Nanopore reads and Illumina reads. The final assembly size is 131.6 Mb, consisting of 765 contigs with a Contig N50 of 1.63 Mb (Table 1). The GC content of the genome assembly is 35.49%. Assembly statistic of *H. hebetor* are compared to four additional braconid genomes, and the result shows a higher N50 value of *H. hebetor* than *Fopius arisanus* (0.98 Mb) [33], *Diachasma alloeum* (0.65 Mb) [23], *Macrocentrus cingulum* (0.19 Mb) [26] and *Microplitis demolitor* (1.1 Mb) [34]. Our analysis indicates that this assembly of a single wasp is more continuous than most wasp genomes which generated by pooled samples.

**Table 1.** Information of the genome assembly.

| Statistic | *Habrobracon hebetor* |
| --- | --- |
| Total length (bp) | 131,644,965 |
| Total length without N (bp) | 131,644,965 |
| Contig number | 765 |
| GC content (%) | 35.49 |
| Contig N50 (bp) | 1,625,734 |
| Contig N90 (bp) | 140,615 |
| Average (bp) | 172,084.92 |
| Median (bp) | 9,847.00 |
| Min (bp) | 177 |
| Max (bp) | 7,063,052 |

### Genome quality assessment

The completeness of the assembly was assessed by using Benchmarking Universal Single-Copy Orthologs (BUSCO) [35], 1,643 out of 1,658 (99%) conserved arthropod genes were found in the genome, 97.3% occurred as single copies (Table 2). The complete and duplicated BUSCO component of the genome was 1.7%. Only three BUSCOs (0.2%) were found fragmented in the genome. There are still twelve BUSCOs (0.8%) cannot be deleted in this genome. We also mapped the Illumina paired-end genomic sequencing reads to the assembled genome, 99.27% of reads could be mapped to the genome. These results indicate that the genome assembly is both highly accurate and near completion.

**Table 2.** BUSCO assessment of the final assembly.

| Category | Number of BUSCOs |
| --- | --- |
| Complete BUSCOs (C) | 1,643 (99.0%) |
| Complete and single-copy BUSCOs (S) | 1,614 (97.3%) |
| Complete and duplicated BUSCOs (D) | 29 (1.7%) |
| Fragmented BUSCOs (F) | 3 (0.2%) |
| Missing BUSCOs (M) | 12 (0.8%) |
| Total BUSCO genes | 1,658 (100%) |

We next mapped *H. hebetor* genome to a genome of *Bracon brevicornis* which is a close relative of *H. hebetor.* The mapping result shows a high-level genome synteny between these two wasps, suggesting the genome assembly of *H. hebetor* obtained from a single wasp is accurate and contiguous (Figure 1). From these results, we didn’t find the evidence to support that amplification biases of WGA could largely influence the quality of genome assembly. We reasoned that might due to the relatively small size genome of parasitoid wasps.

**Figure 1.**
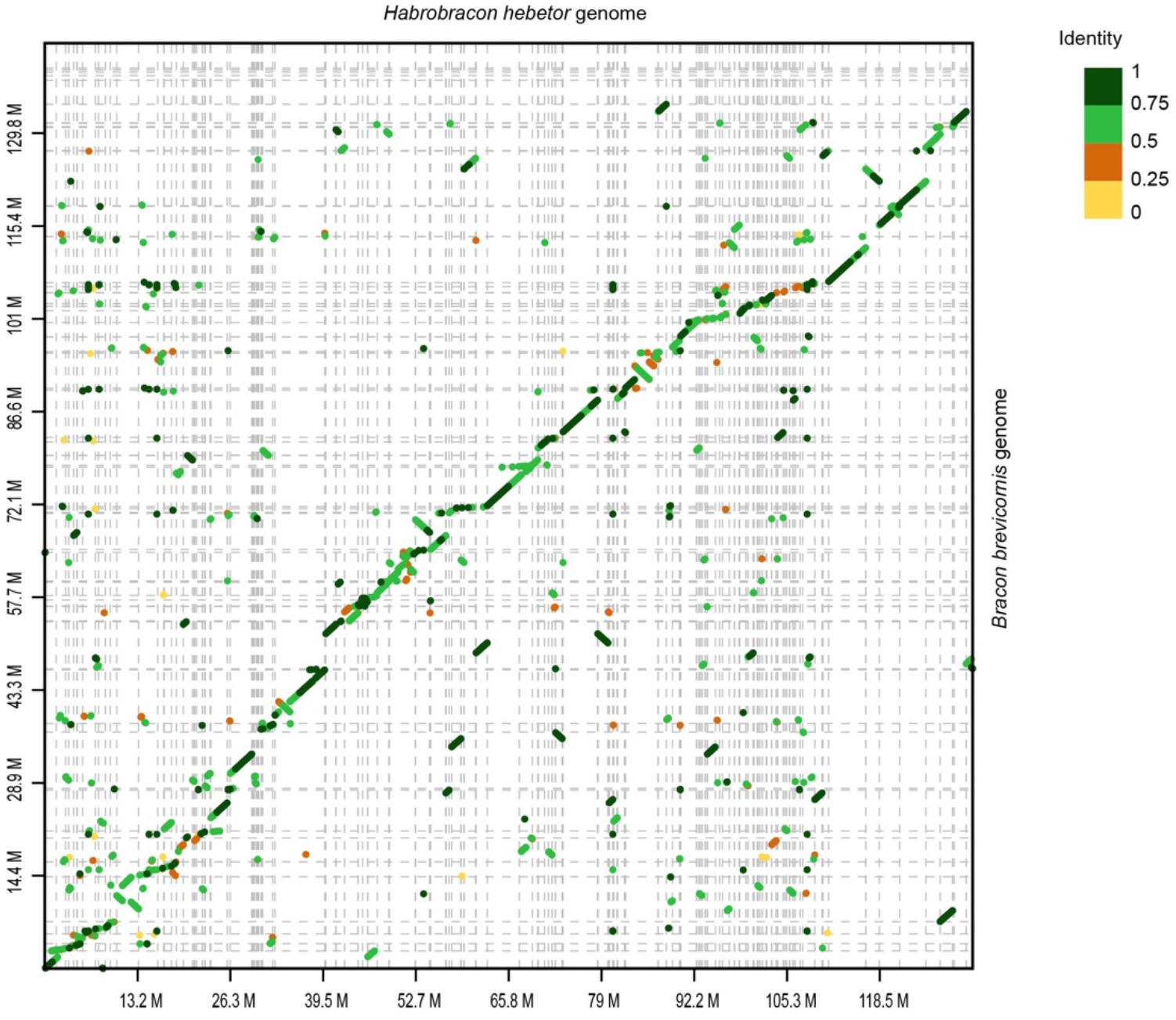
Genome comparison between *H. hebetor* and *B. brevicornis* genome.

In summary, we report a high-quality genome assembly of a single parasitoid wasp *H. hebetor* (~20 ng starting DNA) using WGA, Nanopore and Illumina sequencing technologies. This study presents an example for *de novo* assembling a genome from ultra-low input DNA, which could be used for many small sized species sequencing projects, haploid genomics and population genetics of small sized species.

## Methods

### DNA extraction and whole genome amplification

High molecular weight DNA was extracted from a single male adult *H. hebetor* using TIANamp Micro DNA Kit (DP316) following manufacturer’s recommendations. Two DNA quantification methods Qubit and Nanodrop were used to measure DNA concentration. Then, ~20 ng genomic DNA was amplified using a whole-genome amplification (WGA) kit according to the manufacturer’s instructions (Qiagen REPLI-g Mini Kit, Qiagen, Valencia, CA). The REPLI-g Kit is developed based on multiple displacement amplification (MDA), a WGA method with high processivity and low error rate [36, 37]. Purified genomic DNA was firstly mixed with a denaturation buffer by vortexing and centrifuge briefly. This reaction was quenched by 3 min incubation (at room temperature) with neutralization buffer. The master mix components with REPLI-g Mini DNA Polymerase were added to denatured DNA. The amplification step performed by incubation at 30°C for 16 hours. Next, REPLI-g Mini DNA Polymerase was inactivated by heating the sample for 3 min at 65°C. Following WGA, DNA concentration was determined by using Qubit.

### Nanopore sequencing

A total amount of 2.02 μg DNA was used as input for ONT 1D library construction and sequencing. In brief, the gDNA was sheared using the Megaruptor. Then, the large fragments were selected and purified using AMPure beads. A ONT 1D sequencing library was prepared using the Nanopore Ligation Sequencing Kit (SQK-LSK109; Oxford Nanopore, Oxford, UK) and was sequenced on ONT PromethION 24 platform with one nanopore flow cell (FLO-PRO002).

### Illumina sequencing

Sequencing library was generated using Truseq Nano DNA HT Sample preparation Kit (Illumina USA) following manufacturer’s recommendations. Briefly, the DNA firstly sheared by Covaris S2 system (Covaris, Inc. Woburn, MA, USA), then DNA fragments were end polished, A-tailed, and ligated with the full-length adapter for Illumina sequencing with further PCR amplification. At last, PCR products were purified (AMPure XP system) and libraries were analyzed for size distribution by Agilent2100 Bioanalyzer and quantified using real-time PCR. The final library was sequenced by Illumina NovaSeq platform.

### Genome assembly

Nanopore long reads flagged as ‘‘passing’’ were corrected by NECAT (https://github.com/xiaochuanle/NECAT). Flye (version: 2.7.1-b1590) [31, 32] was used to assemble the genome with default parameters using >5K Nanopore reads (~250X). Then, Racon (https://github.com/isovic/racon) was used for correcting the assembly. In addition, iterative polishing was conducted using Pilon (version: 1.22) [38] with adapter-trimmed paired-end Illumina reads. The Pilon program was run with default parameters to fix bases, fill gaps, and correct local misassemblies.

### Evaluation

Benchmarking Universal Single-Copy Orthologs method (BUSCO version 4.0) [35] was used to search the 1,658 bench-marking universal single-copy orthologous genes in insecta_odb9.

### Genome comparison

An online tool D-GENIES [39] was used to compare the *H. hebetor* and *B. brevicornis* genome [40].

## Data availability

All sequence data are available at the NCBI, Bioproject number PRJNA644201.

## Acknowledgements

This work was supported by Key Program of National Natural Science Foundation of China (NSFC) (Grant no. 31830074 to GYY), Major International (Regional) Joint Research Project of NSFC (Grant no. 31620103915 to GYY), XHY and LX thank “Academic Star” Program for Ph. D Student of Zhejiang University for support. XHY thanks the China Scholarship Council (No. 201906320376) for its support during his research period in the Rochester, New York, US. We thank Prof. John H. Werren (University of Rochester) for valuable discussions.

## Author contributions

GYY conceived and designed the works, and supervised the project. GYY, FL and QF coordinated the project. KLY and SJX prepared the samples for sequencing. XHY, ZYT, HC sequenced and assembled the genome. XHY and YY performed the bioinformatics analysis. XHY wrote the draft manuscript. YY, LX, SX, CLY, HC, FL, QF and GYY improved and revised the manuscript. All authors read and approved the final manuscript.

---

**Table S1.** Nanopore sequencing data.

| Type | TotalBase | TotalReads | MaxLen | AvgLen | N50 | N90 | meanQ |
| --- | --- | --- | --- | --- | --- | --- | --- |
| >0 | 51,597,252,675 | 11,507,945 | 124,237 | 4,484 | 7,311 | 2,127 | 11 |
| >2000 | 46,957,240,951 | 7,372,598 | 124,237 | 6,369 | 8,039 | 3,113 | 11 |
| >5000 | 34,318,006,006 | 3,510,066 | 124,237 | 9,777 | 10,379 | 5,877 | 11 |
| >10000 | 18,071,218,227 | 1,193,518 | 124,237 | 15,141 | 14,941 | 10,769 | 11 |
| >20000 | 4,376,715,930 | 170,092 | 124,237 | 25,731 | 24,876 | 20,737 | 11 |
| >100000 | 534,472 | 5 | 124,237 | 106,894 | 102,023 | 100,525 | 10 |

**Table S2.** Illumina sequencing data.

| Raw Reads | Clean Reads | Clean Base (Gb) | Q20(%) | Q30(%) | GC Content (%) |
| --- | --- | --- | --- | --- | --- |
| 46,619,030 | 46,061,685 | 13.82 | 95.51 | 91.23 | 35.27 |

**Figure S1.**
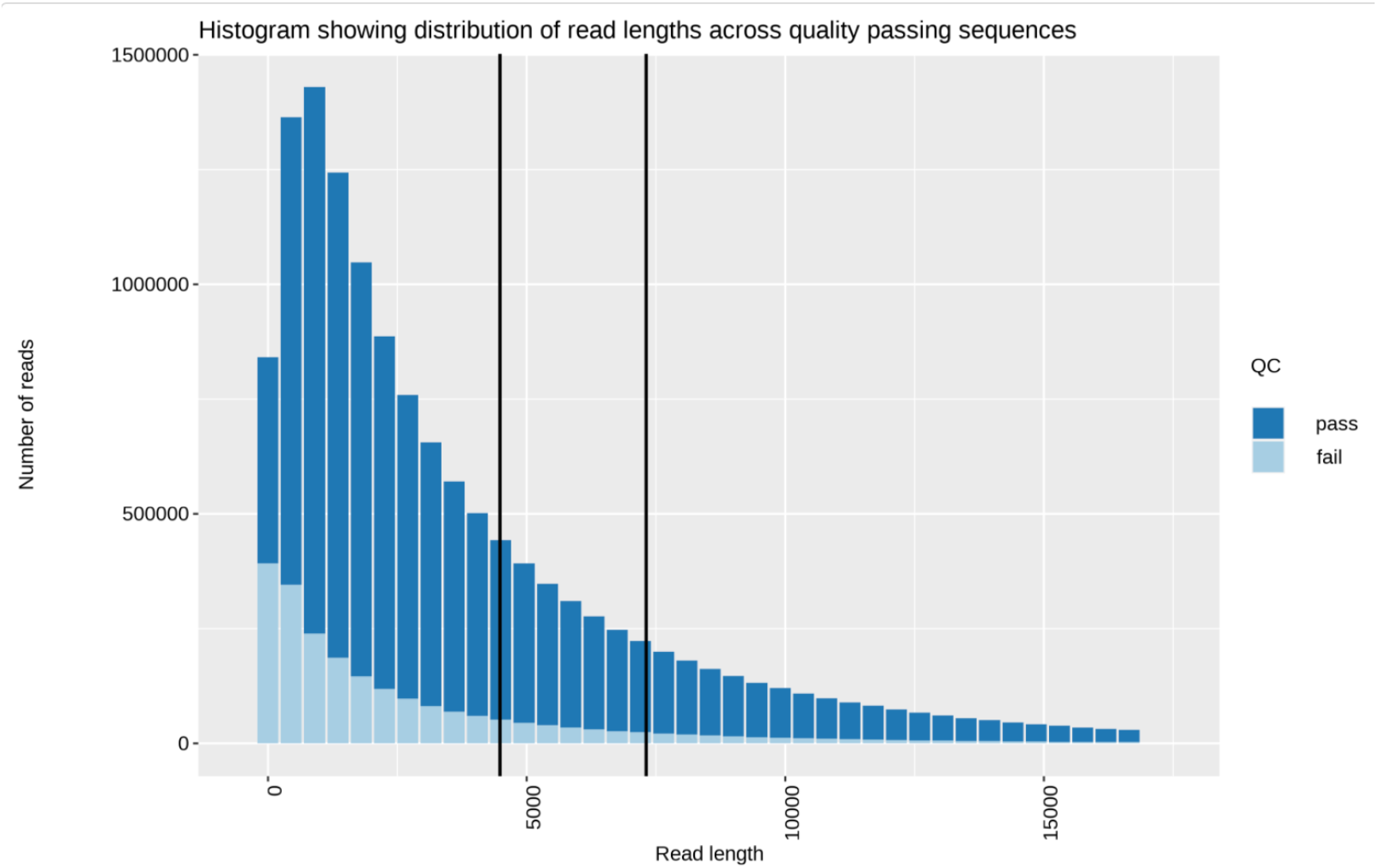
Distribution of the Nanopore reads.

